# 3-D Vascularized Breast Cancer Model to Study the Role of Osteoblast in Formation of a Pre-Metastatic Niche

**DOI:** 10.1101/2021.05.05.442719

**Authors:** Rahul Rimal, Prachi Desai, Andrea Bonnin Marquez, Karina Sieg, Yvonne Marquardt, Smriti Singh

**Affiliations:** DWI - Leibniz Institute for Interactive Materials, Forkenbeckstrasse 50, 52074 Aachen, Germany; Department of Dermatology and Allergology, University Hospital, RWTH Aachen University, 52074 Aachen, Germany

**Author notes:** Dr. S. Singh, Max-Planck-Institut für medizinische Forschung, Jahnstraße 29, 69120 Heidelberg, Germany.

**Keywords:** breast cancer, triple negative, tumor-microenvironment, pre-clinical, bone metastasis, blood vessels, scaffold-free models

## Abstract

Breast cancer cells (BCCs) preferentially metastasize to bone. It is known that BCCs remotely primes the distant bone site prior to metastasis. However, the reciprocal influence of bone cells on the primary tumor is relatively overlooked. Here, to study the bone-tumor paracrine influence, a tri-cellular 3-D vascularized breast cancer tissue (VBCTs) model is engineered which comprised MDA-MB231, a triple-negative breast cancer cells (TNBC), fibroblasts, and endothelial cells. The VBCTs are indirectly co-cultured with osteoblasts (OBs), thereby constituting a complex quad-cellular tumor progression model. MDA-MB231 alone and in conjunction with OBs led to abnormal vasculature and reduced vessel density but enhanced VEGF production. A total of 1476 significantly upregulated and 775 downregulated genes are identified in the VBCTs exposed to OBs. HSP90N, CYCS, RPS27A, and EGFR are recognized as upregulated hub-genes. Kaplan Meier plot shows HSP90N to have a significant outcome in TNBC patient survivability. Furthermore, compared to cancer tissues without vessels, gene analysis recognized 1278 significantly upregulated and 566 downregulated genes in VBCTs. DKK1, CXCL13, C3 protein and BMP4 are identified to be downregulated hub genes in VBCTs. Together, a multi-cellular breast cancer model and culture protocols are established to study pre-metastatic events in the presence of OBs.

## 1. Introduction

Breast cancer is the most frequently diagnosed cancer and the leading cause of cancer-related death in females worldwide.^1^ Once the tumor metastasizes to a distant site, the 5-year survival chance decreases from ~ 93% to ~ 22% (National Cancer Institute SEER database). There are five molecular subtypes of breast cancer of which the triple negative subtype is particularly challenging.^2^ Triple negative tumors represent around 15% of invasive ductal breast cancers. They display distinctive epidemiological, phenotypic and molecular features with distinctive patterns of relapse, and a poor prognosis despite relative chemosensitivity.^2^ Increasing clinical evidence has suggested that the most common site of triple negative breast cancer (TNBC) metastasis is bone,^3–6^ which disrupts the balance between bone formation and resorption.^7^ Patients with this condition have a median survival of around 2-3 years following initial diagnosis of bone involvement.

To understand better the propensity of breast cancer cells (BCCs) to metastasize preferentially to bone, an investigation of the mechanistic pathways that drive organ-specific metastasis is crucial. Conventionally, researchers have relied on two-dimensional (2-D) assays and animal models in order to study the metastatic process.^8^ However, 2-D models do not recapitulate the tissue architecture and the cellular interactions of the tumor microenvironment. On the other hand, animal models have failed to translate pre-clinical success to patient outcome, thereby questioning its resemblance in the cellular level to the human tissue. With animal models, it is difficult to study the interaction of specific cell types in a controlled manner thereby limiting the possibility to conduct parametric studies. Additionally, ethical issues in using animals as models for human disease exist. An alternative approach is the use of three-dimensional (3-D) cultures such as hydrogels composed of either ECM molecules^9–11^ or synthetic polymers,^12^ scaffold-free organoids culture,^13, 14^ and microfluidic chips.^15, 16^

Although there are a large number of prior studies focused on the fabrication of sophisticated 3-D *in-vitro* models utilizing the above-mentioned techniques, most models are not developed for the investigation of cancer metastasis progression. Out of the limited number of research articles that deal with breast-to-bone metastasis, the majority of these studies recapitulate the late-bone metastatic stage where the BCCs have already colonized the bone microenvironment, leading to further metastasis or dormancy.^8, 17, 18^ In other words, these studies focus on post-metastatic, direct interaction between BCCs and bone. However, to understand the metastasis process in its entirety, the study of early-stage pre-metastatic progression of BCCs within the primary site is equally important.

One overlooked pre-metastatic phenomenon is the involvement of the distant secondary site in remotely regulating the primary site. Bone could act as an active participant, releasing cytokines and growth factors to the primary breast tumor site, and play a functional role in the initiation of tumor metastasis.^19^ The concept that primary tumors can prime the secondary site remotely prior to the initiation of metastasis is not controversial.^20, 21^ However, the reciprocal crosstalk from the secondary site, although understudied, could also influence changes in the primary site. It is intriguing to note that high bone mineral density (BMD) in postmenopausal women has been linked with an increased incidence rate of breast cancer.^22, 23^ Furthermore, it was shown that low levels of bone turnover proteins c-terminal telopeptide (CTX1, bone resorption marker) and osteocalcin (bone formation marker), independent of BMD, correlated with increased breast cancer risk in postmenopausal women.^24^ Although these phenomena are not fully understood and require further investigation, it begs the question of whether the bone can indirectly, via release of soluble factors, lead to breast cancer initiation and progression.^19^

To mimic this early-stage pre-metastatic cross-talk scenario *in-vitro*, there is a need to develop a 3-D indirect culture protocol where the secondary and primary sites are not in direct contact but interact via indirect paracrine signaling. Due to the lack of such models and protocols, the study of organ specificity (organotropism) observed in metastasis remains a challenge and several open questions like the role of future secondary sites in conditioning the primary tumor persists. The goals of our study were therefore to engineer a 3-D vascularized breast cancer tissue (VBCTs) representing the primary tumor site indirectly co-cultured together with OBs, representing the secondary site. We further strived to study whether and how this indirect interaction with bone cells could lead to alterations in the primary breast tumor microenvironment. Therefore, a quad-culture system was developed consisting of TNBCs (MDA-MB231), fibroblasts, and endothelial cells (HUVECs) for indirect co-culture with OBs, the bone-forming cells. We opted for a scaffold-free model system due to its several advantages over scaffold-based models.^25^ To engineer the scaffold-free models, we utilized layer-by-layer ECM coating and accumulation technique.^26^ This technique allowed the formation of thick vascularized tissues and provided a microenvironment to the cancer cells with enhanced direct cell-cell interactions.^26, 27^ Fabrication of the models in cell-culture inserts allowed easy handling and the indirect interaction of the formed VBCTs (within the insert) with the OBs (on the well plate).

We show the formation of a vascularized breast cancer model using MDA-MB231 and the influence of OBs on the VBCT morphology, vasculature and gene expression. Furthermore, the effect of blood vessels (BVs) on the model in presence of OBs was elucidated. We observed that the morphology of MDA-MB231 changes significantly in 3-D scaffold-free tissues as compared to 2-D cultures. Following this, we investigated the influence of OBs and MDA-MB231 on the tissue vasculature. Finally, utilizing gene microarray analysis, the indirect paracrine effect of OBs on the VBCTs was studied. Since vascular re-modeling is one of the first steps in the initiation of a pre-metastatic niche, we also studied the influence of BVs in the breast cancer tissues co-cultured with OBs. Important hub genes and highly interconnected protein-protein interaction (PPI) clusters were identified to understand the influence of OBs and BVs in the VBCTs.

In short, we fabricated a simple and realistic biological system that was utilized to understand the influence of the secondary bone site on the primary breast cancer microenvironment. This system, in the future, could be used to dissect the role of secondary tumors that drives breast cancer organotropism.

## 2. Results

### 2.1 Formation of 3-D vascularized breast cancer tissues

To fabricate the scaffold-free breast cancer model, we utilized the layer-by-layer ECM coating and accumulation technique. Normal fibroblasts (NFs) were coated with fibronectin (FN) and gelatin (G) proteins to form nano-layer of ECM moieties around the cells.^20, 28^ The coated individual NFs were mixed with HUVECs and MDA-MB231 BCCs followed by accumulation in a cell-culture insert. We observed that MDA-MB231 morphology was altered in 3-D scaffold-free tissues as compared to 2-D culture **(Figure 1a, b, e)**. We observed that the BCCs interacted with the BVs either by extending along the vascular length or by remaining in a circular morphology in direct contact with the BVs **(Figure 1c, d)**. Majority of the BCCs showed an extended morphology and elongated filopodia in the vascularized 3-D culture **(Figure 1b, c, e)**. Similar filopodia structures in MDA-MB231 BCCs were observed previously in a 3-D collagen matrix.^29^ We quantified the BCCs aspect ratio and circularity and noted that the BCCs morphology in 3-D ranged from low (circular) to high (elongated) while BCCs in 2-D flat surface did not demonstrate elongated morphology **(Figure 1f, g).**

**Figure 1.**
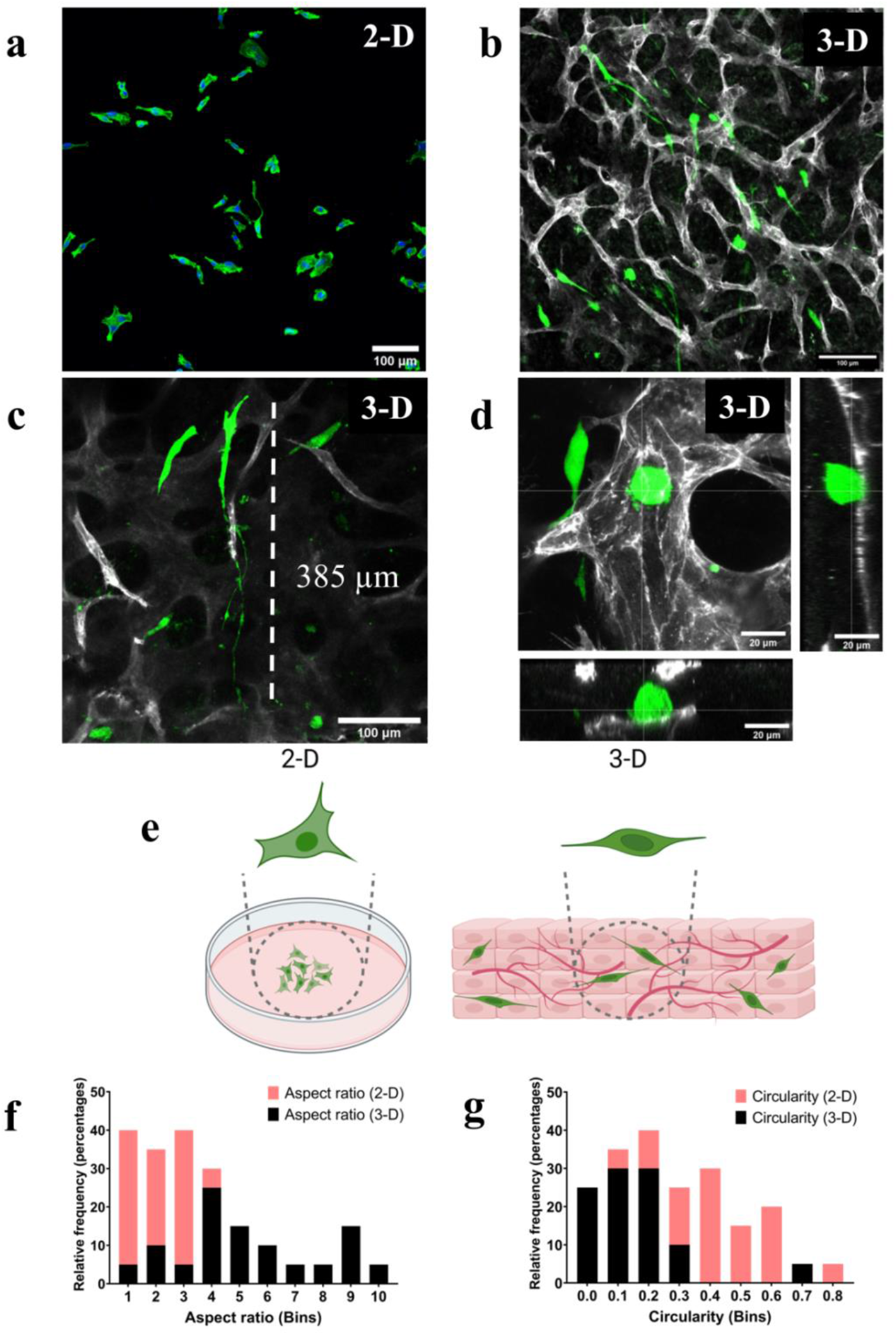
Formation of *in-vitro* vascularized breast cancer tissue. a) Morphology of GFP^+^ MDA-MB231 (green) in 2-D cell culture plate, Scale=100 μm b) morphology of GFP^+^ MDA-MB231 embedded in scaffold-free vascularized tissue, vessels were stained (grey) with PECAM1 (CD-31), Scale=100 μm c) highly elongated BCCs filopodia observed in 3-D tissues wrapped around a vessel branch, Scale=100 μm d) direct interaction between cancer cells and BVs, orthogonal projections are shown e) Pictorial representation of the change of morphology of MDA-MB231 in 2-D culture and 3-D scaffold-free culture f) cell aspect ratio in 2-D is in the lower range as compared to cells in the 3-D models that range from low to high (data from randomly selected 20 cells from 3 samples) g) cell circularity (1=circular, 0=elongated) analysis show that cells in 3-D tend to be more elongated as compared to 2-D cultures (randomly selected 20 cells from 3 samples).

Recently, it was shown that MDA-MB231 morphology fluctuated depending on the ECM composition of the substrate.^30^ The presence of FN led to an increase in the formation of tunneling nanotubes, whereas the presence of matrigel led to highly elongated cell morphology.^30^ Furthermore, MDA-MB231 morphology has been shown to change drastically in collagen substrates of varying concentrations suggesting that the morphology is highly dependent on its surrounding microenvironment.^30^

Since our model is comprised of cell-secreted ECM rather than preferentially pre-selected ECM hydrogels with designated concentration, we believe that the scaffold-free models depict an unbiased tumor-microenvironment and therefore predict better the *in-vivo* cellular morphology.

### 2.2 Presence of MDA-MB231 and OBs modulates vasculature

Firstly, to confirm that the OBs seeded on the well plate **(Figure S1f)** can influence the BCCs within the cell-culture insert, we conducted a 2-D migration assay to observe the migration of BCCs due to OBs. We noted an enhanced migration of MDA-MB231 cells in the presence of OBs or 100ng/ml C-X-C-chemokine ligand 12 (CXCL12) **(Figure S1a-g)**. Previous studies have shown that the positive-feedback loop initiated by the C-X-C chemokine receptor type 4 (CXCR4), highly expressed in malignant BCCs, to the CXCL12 expressed in bone lead to breast cancer migration.^31,32^ We noted that the pre-treatment of BCCs with CXCR4 antagonist AMD3100 (25μg/ml) restricted this migration of BCCs towards both OBs and CXCL12 **(Figure S1d, e, g)**. Therefore, we confirmed that cell culture inserts could be utilized for indirect co-culture assays where the secreted factors of one cell could influence the other cell type.

Following this, we fabricated the VBCTs within the insert and seeded OBs on the bottom of the well plate to study the influence of OBs on vascularization of the tissue. A schematic illustration of the setup is shown in **Figure 2a**. For testing the effect of BCCs and OBs on vessel fate, four conditions were tested namely vascularized non-cancerous tissue (NH), vascularized tissue with MDA-MB231 (NHM), vascularized non-cancerous tissue indirectly co-cultured with OBs (NHO) and, vascularized tissues with MDA-MB231 indirectly co-cultured with OBs (NHMO). For clarity, the conditions and their cell constituents are summarized in **Table 1**.

**Figure 2.**
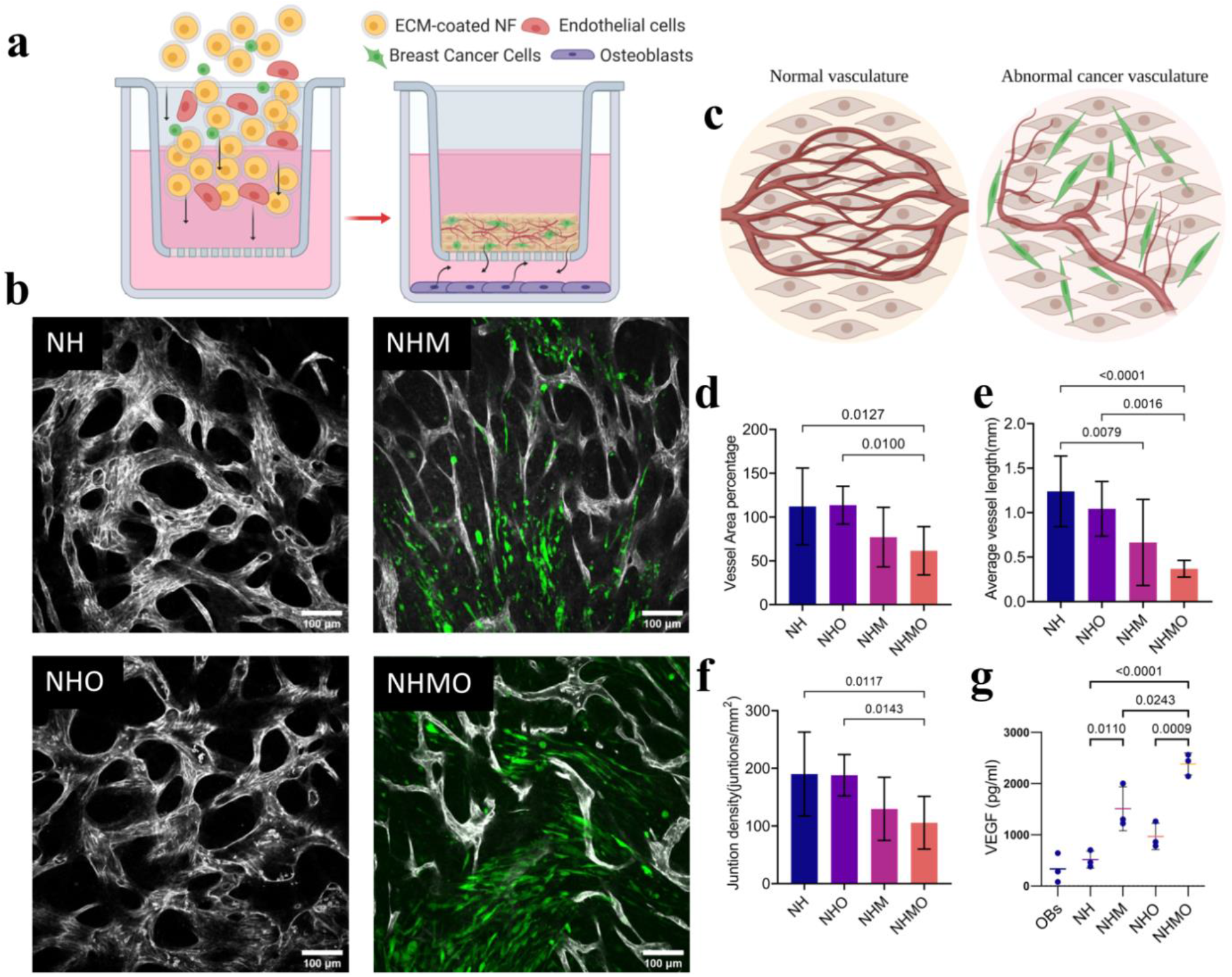
Influence of BCCs and OBs on blood vessel architecture. a) Schematic overview of the indirect co-culture experiment and the interaction of several cell types b) Confocal images of tissues (BVs morphology was monitored after day 7 of culture). BVs were observed in NH, NHM, NHO, and NHMO tissue samples. Vessels were stained with PECAM-1 (grey) and MDA-MB231 (GFP^+^, green). Scale bar=100 μm. c) Schematics show abnormal changes of BVs in the presence of MDA-MB231. OBs accentuated the abnormality of the BVs d) Analysis shows a significant decrease in vessel area percentage in NHMO samples as compared to NH and NHO samples (n=3, with 3 different tissue area per sample) e) significantly decreased average vessel length (mm) in NHMO as compared to NH and NHO, and significantly decreased vessel length in NHM as compared to NH, (n=3, with 3 different tissue area per sample), f) significantly decreased junction density in NHMO as compared to NH and NHO, (n=3, with 3 different tissue area per sample) g) VEGF analysis shows a significant difference in VEGF levels between NH and NHM, NHO and NHMO, and NH and NHMO, VEGF level in OBs monolayer alone was statistically different to NHM, NHO and NHMO but not significantly different to NH (p-values not shown for graph simplicity), (n=3). Values in all the graphs are mean ± standard deviation; analysis was done using ordinary one-way Anova.

**Table 1.** List of different tissue conditions.

| Abbreviation | Fibroblasts<br>(N) | HUVECs<br>(H) | MDA-MB231<br>(M) | Osteoblasts<br>(O) |
| --- | --- | --- | --- | --- |
| NH | + | + | - | - |
| NHM | + | + | + | - |
| NHO | + | + | - | + |
| NHMO | + | + | + | + |
| NMO | + | - | + | + |
| OBs | - | - | - | + |

We observed that the NH and NHO tissues formed an organized vascular network, whereas NHM and NHMO models showed abnormal vascularization with disorganized architecture **(Figure 2b)**. A pictorial representation is shown in **Figure 2c**. We noted that the vessel area percentage and junction density was significantly lower in NHMO samples as compared to NH and NHO samples **(Figure 2d, f)**. Similarly, the average vessel length was significantly lower in NHMO as compared to NH and NHO **(Figure 2e)**. The average vessel length was also lower in NHM as compared to NH **(Figure 2e)**. These readings indicate that in a scaffold-free system, the inclusion of BCCs leads to detrimental alterations in the surrounding vasculature and this process is accentuated when the cancerous tissue is exposed indirectly to OBs.

To further understand the influence of OBs and BCCs derived soluble factors on angiogenesis, we conducted the VEGF-ELISA on day 7 of culture time. Previous reports have indicated that the VEGF levels in the intra-tumoral region of TNBC tissues were significantly higher than non-TNBC tissues.^33^ Elevated VEGF level in breast cancer patients has also been linked to diminished progression-free survival and overall survival of the patient.^34^ OBs have also been shown to be a source of VEGF, particularly during bone repair.^35^ We observed the highest VEGF secretion in NHMO condition followed by NHM, NHO, NH and OBs respectively. VEGF levels from NHMO were significantly different to those from NHM, NH, NHO, NH and OBs **(Figure 2g)**. VEGF levels from NHM were significantly different to NH and OBs while NHO was significantly different to OBs **(Figure 2g)**.

We can therefore conclude that the presence of MDA-MB231 in the *in-vitro* tissue led to increased production of VEGF, and the indirect co-culture with OBs elevates the VEGF levels.

In the present cancer model, the vessel area percentage or junctional density in the vascularized tissue did not correlate with the increase in VEGF levels. Instead, we observed segmented, constricted and abnormal BVs in the presence of MDA-MB231 that led to the decrease in vascular density.

The current cancer model comprised BCCs distributed randomly within the 3-D tissue. To better deliberate the interaction of BCCs and the vasculature, we fabricated a separate model where cancer cells were seeded on top of the vascularized tissue construct instead of being randomly distributed **(Figure S2a)**. As the MDA-MB231 cells were grouped in a single layer, it was expected that the direct interaction of BCCs with the vascular bed beneath would better demonstrate the fate of vessels. At the intersection between cancer cells and vasculature, we observed bright PECAM-1^+^ spots that indicated broken or open vascular ends **(Figure S2c)**. Magnified images of the spots showed that the open vascular ends have thin irregular sprouts protruding out of them **(Figure S2b, d)**.

### 2.3 OBs modulates gene expression in VBCTs

Our next focus was to elucidate the influence of OBs on the tumor tissue. VBCTs were cultured indirectly with OBs for 7 days and subsequently harvested. Gene microarray analysis was conducted to identify differentially expressed genes (DEGs) and crucial hub genes. We analyzed the DEGs between NHM and NHMO samples to elucidate the influence of OBs on the VBCTs. Upregulated genes (Fold change, FC ≥2, p<0.05) and downregulated genes (FC ≤−2, p<0.05) were selected as the significant DEGs.

We observed 1476 significantly upregulated genes and 775 significantly downregulated genes in models that were co-cultured with OBs. The volcano plot depicts the upregulated and downregulated genes **(Figure 3a)**. The top five upregulated DEGs and the top five downregulated DEGs are shown in **Table 2**. Each of the highly upregulated genes was loaded into TNM plotter (BRCA database) to observe the differences in their expression in normal, cancerous and metastatic tissues **(Figure S4a-e)**.^36^ We observed that all the top five upregulated genes in NHMO samples as compared to NHM samples were expressed more in cancerous and metastatic breast cancer tissues as compared to normal breast tissues **(Figure S4a-e)**.

**Figure 3.**
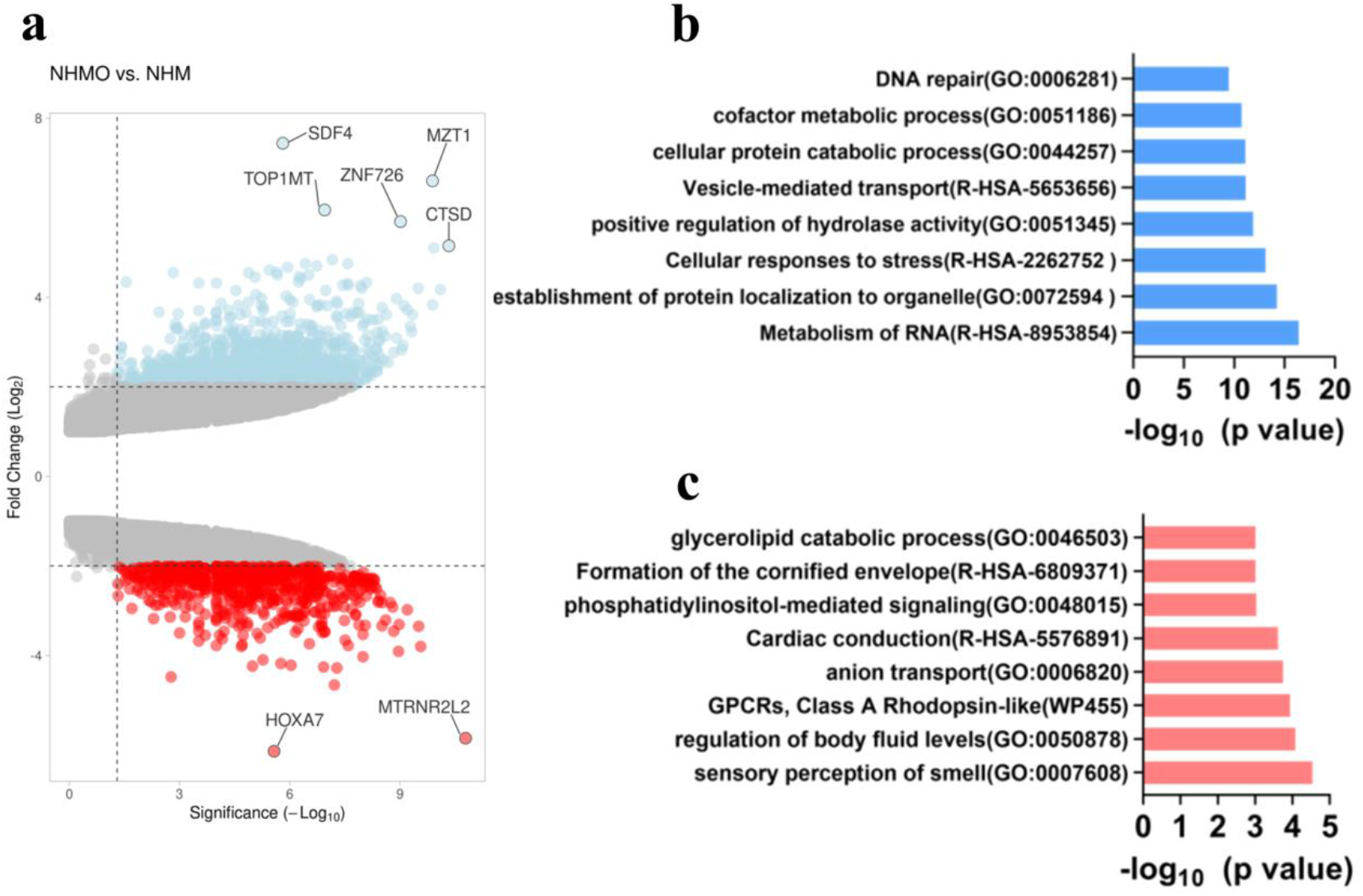
a) volcano plot depicts differentially expressed genes in NHMO vs NHM (red=downregulated, blue=upregulated) b) top upregulated gene ontology terms c) top downregulated gene ontology terms

**Table 2.** Top five upregulated and downregulated genes (NHMO vs. NHM)

| Gene symbol | Gene name | Expression | FC | P-value |
| --- | --- | --- | --- | --- |
| *SDF-4* | stromal cell derived factor 4 | up | 7.44 | 1.55E-06 |
| *MZT1* | mitotic spindle organizing protein 1 | up | 6.6 | 1.31E-10 |
| *TOP1MT* | topoisomerase (DNA) I, mitochondrial | up | 5.95 | 1.13E-07 |
| *ZNF726* | zinc finger Protein 726 | up | 5.69 | 9.70E-10 |
| *CTSD* | cathepsin D | up | 5.15 | 4.70E-11 |
| *HOXA7* | homeobox A7 | down | -6.14 | 2.68E-06 |
| *MTRNR2L2* | MT-RNR2-like 2 | down | -5.85 | 1.65E-11 |
| *MAGEA2* | MAGE family member A2 | down | -4.66 | 6.13E-08 |
| *A2BP1* | Ataxin-2 binding protein 1 | down | -4.48 | 0.0017 |
| *PDZK1IP1* | PDZK1 interacting protein 1 | down | -4.28 | 5.23E-08 |

The gene signature consisting of all the five upregulated genes was observed to be expressed more in both TNBC and broad BRCA tissue database (fed into GEPIA2 application) as compared to normal tissue database **(Figure S4f)**. The functional enrichment analysis was conducted on the upregulated and downregulated DEGs separately **(Figure 3b, c)**. Five top upregulated pathways were RNA metabolism, the establishment of protein localization to organelle, cellular response to stress, positive regulation of hydrolase activity and vesicle mediated transport **(Figure 3b)**. Five top downregulated pathways were sensory perception of smell, regulation of body fluid levels, GPCRs-Class A Rhodopsin like anion transport and cardiac conduction **(Figure 3c).**

Next, the four highly interconnected hub genes from the upregulated PPI network were identified namely HSP90N, CYCS, EGFR, and RPS27A. The expression of the four hub genes was tested in tumor and normal tissue database (fed into GEPIA2 application) specific for TNBC patients (**Figure 4 a-d**). The result from the TNBC database indicated that HSP90N and CYCS are significantly present in the tumor tissue as compared to normal breast tissue (**Figure 4 a, c**), and EGFR is expressed significantly more in normal breast tissue as compared to breast cancer tissues (**Figure 4 b**). Disease-free survival plots of the hub genes specific for TNBC patients are shown (**Figure 4 e-h**). Out of the four hub genes, the survival plots indicated that HSP90N had a significantly unfavorable impact on the survival of the patients (**Figure 4g**).

**Figure 4.**
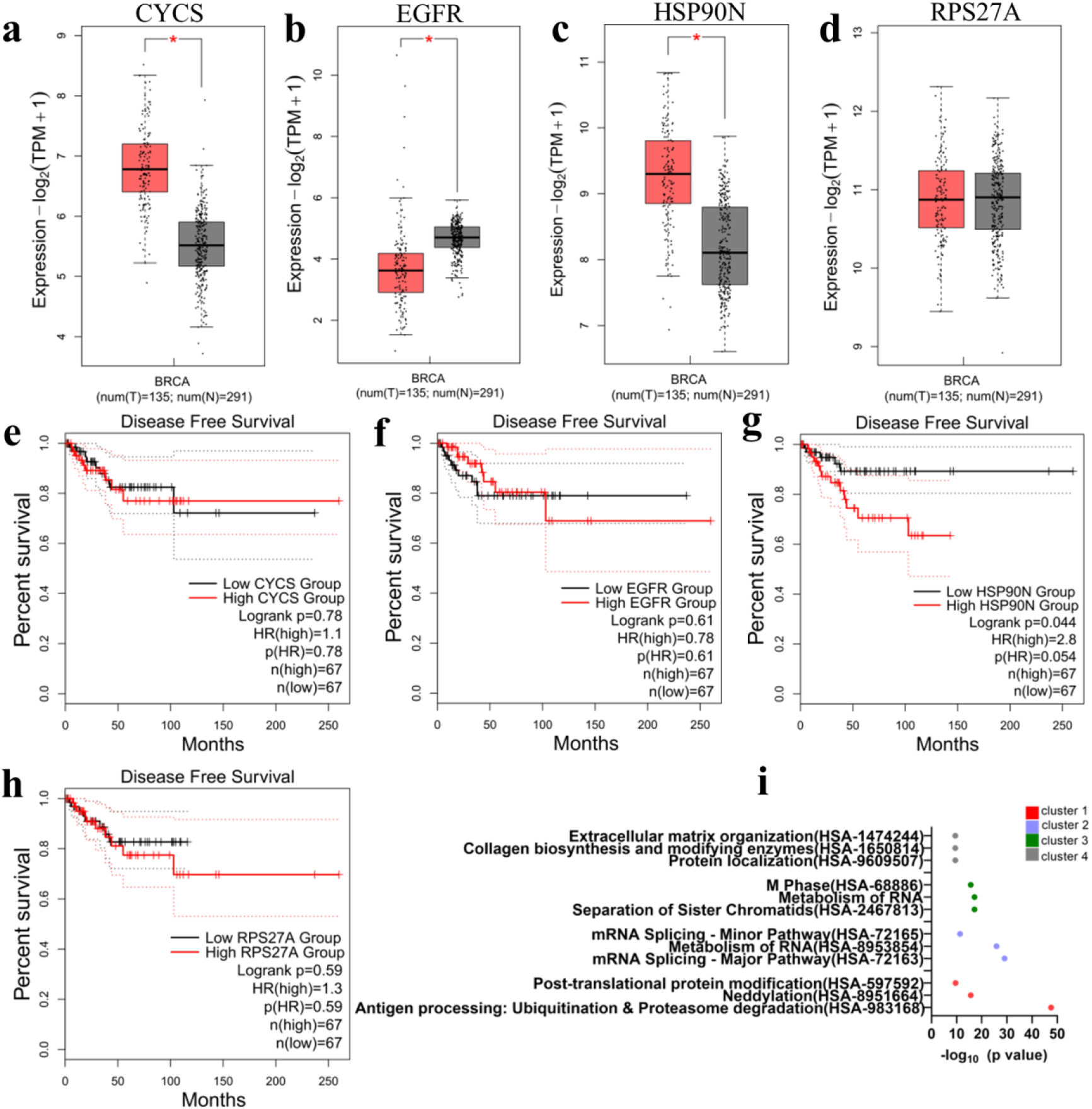
Four upregulated hub genes CYCS, EGFR, HSP90N and RPSP27A were analyzed for their impact on patient outcome. a, b, c, d) Presence of CYCS, EGFR, HSP90N and RPS27A in tumor (red) vs normal (gray) tissue respectively (significance<0.05) e, f, g, h) Disease-free survival plots of CYCS, EGFR, HSP90N and RPS27A respectively i) Reactome enrichment of the clusters of upregulated genes shows four highest clusters recognized (cluster 1-4).

Next, four clusters of highly interconnected upregulated genes were determined **(Figure S5).** Reactome pathway enrichment analysis of each cluster was noted **(Figure 4i)**. The pathways included terms such as antigen processing, neddylation, metabolism of RNA, collagen biosynthesis and modifying enzymes, and ECM organization. **Table S1** shows a complete list of genes involved in each highly interconnected cluster.

### 2.4 Endothelial cells modulate gene expression in breast cancer tissues

We sought to understand the importance of BVs in the breast cancer tissues (in the presence of OBs). Previous reports have suggested that BVs are not merely passive onlookers but active players in the metastasis process.^37^ The comparison between cancerous tissues with vessels, and cancerous tissues without vessels could elucidate the influence of BVs in cancer progression. Therefore, we compared the gene expression between day 7 NHMO and NMO samples.

We observed 1278 upregulated genes and 566 downregulated genes in the presence of BVs. The volcano plot depicting the upregulated and downregulated genes is shown in **Figure 5a**. **Table 3** shows the list of the top five most highly upregulated and downregulated genes.

**Figure 5.**
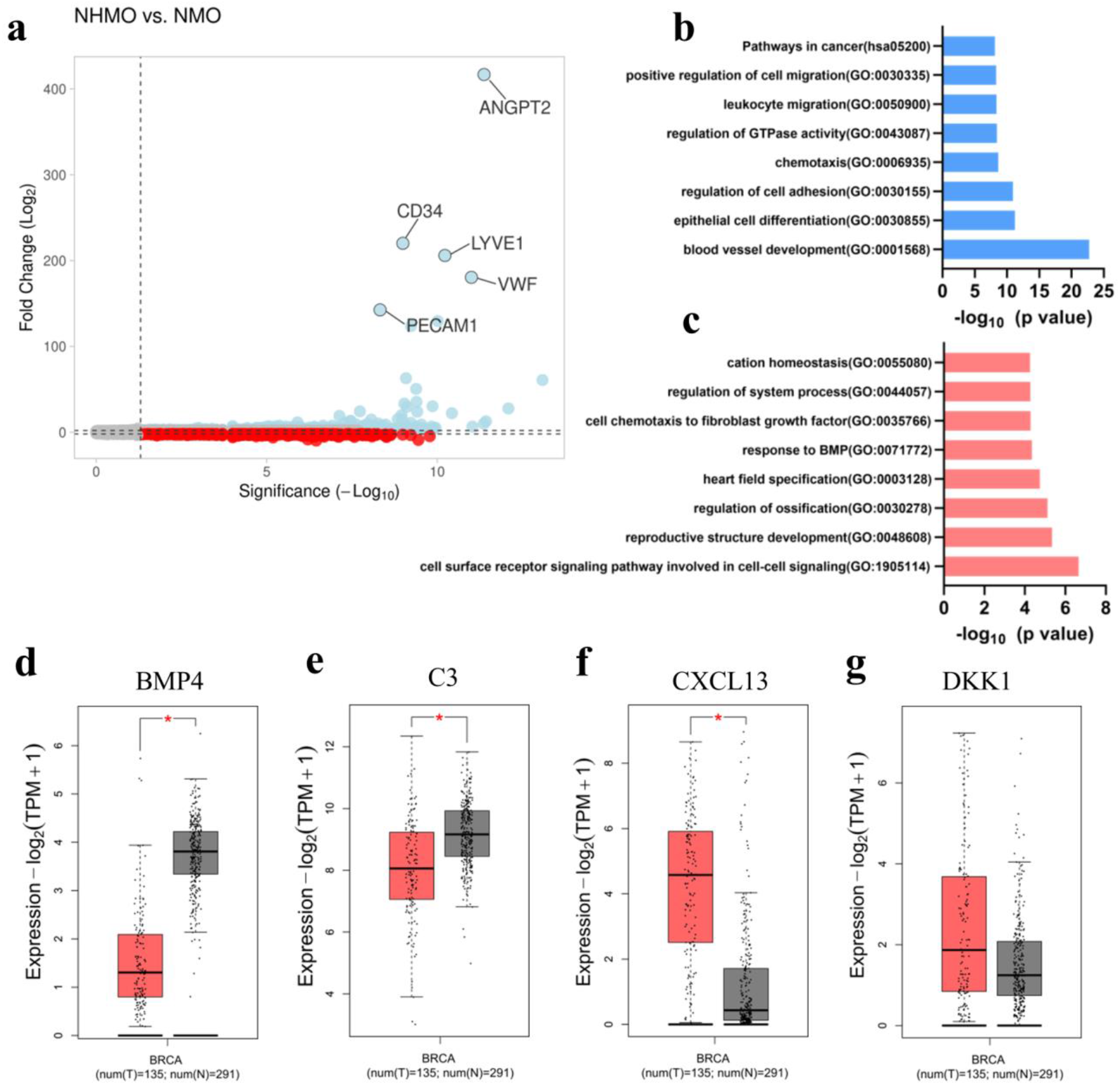
a) volcano plot shows upregulated (blue) and downregulated (red) genes. The top five highly expressed genes are labeled b) Upregulated gene ontology terms c) Downregulated gene ontology terms d, e, f, g) Expression of downregulated hub genes BMP4, C3 protein, CXCL13, and DKK1 (red=tumor tissue, gray= normal tissue), significance <0.05.

**Table 3.** Top five upregulated and downregulated genes due to the presence of BVs (NHMO vs. NMO)

| Gene symbol | Gene name | Expression | FC | P-value |
| --- | --- | --- | --- | --- |
| *ANGPT2* | angiopoietin 2 | up | 416.87 | 4.25E-12 |
| *CD34* | CD34 molecule | up | 220.18 | 1.02E-09 |
| *LYVE1* | lymphatic vessel endothelial hyaluronan receptor 1 | up | 205.97 | 5.98E-11 |
| *VWF* | von Willebrand factor | up | 180.5 | 9.97E-12 |
| *PECAM1* | platelet/endothelial cell adhesion molecule 1 | up | 142.58 | 4.75E-09 |
| *SMOC2* | SPARC related modular calcium binding 2 | down | -5.83 | 9.98E-05 |
| *ADGRD1* | adhesion G protein-coupled receptor D1 | down | -6.1 | 6.57E-06 |
| *CXCL14* | chemokine (C-X-C motif) ligand 14 | down | -8.35 | 6.24E-07 |
| *PTGDS* | prostaglandin D2 synthase 21kDa (brain) | down | -8.95 | 3.58E-10 |
| *PCSK5* | proprotein convertase subtilisin/kexin type 5 | down | -9.67 | 3.55E-07 |

The upregulation of endothelial-related genes was expected as one tissue condition consisted of BVs and the other condition was non-vascularized. The ECs-related genes such as ANGPT2, CD34, LYVE1, VWF and PECAM1 were highly upregulated in the vascularized tumor tissue as compared to non-vascularized tissue due to the presence of BVs **(Table 3)**.

Interestingly, several of the downregulated genes in NHMO vs NMO may have implications in OBs fate and breast cancer progression. The five highly downregulated genes were PCSK5, PTGDS, CXCL14, ADGRD1, and SMOC2 **(Table 3)**. SMOC2, a calcium-binding protein is known to inhibit osteogenesis and prevent mineralization in OBs.^38^ ADGRD1 also known as GPR133 could also have implications in bone diseases.^39^ CXCL14 has shown to be expressed in the stromal cells in malignant breast cancer^40^ and has been shown to induce lung cancer metastasis to bone.^41^

Next, the top functional enrichment pathways of the upregulated genes were determined namely blood vessel development, epithelial cell differentiation, regulation of cell adhesion, chemotaxis, regulation of GTPase activity, leukocyte migration, positive regulation of cell migration and pathways in cancer **(Figure 5b)**. The genes involved in these upregulated pathways are shown in **Table S2**. Two highly interconnected gene clusters are determined from the upregulated PPI network in **Figure S6**.

The downregulated pathways were cell surface receptor signaling pathway, reproductive structure development, regulation of ossification, heart field specification and response to BMP **(Figure 5c)**. The genes involved in these downregulated pathways are shown in **Table S3**. Four hub genes from the upregulated PPI network were identified namely KDR, ITGAM, CXCR4, and PECAM1. Since the upregulated hub genes were mostly associated with endothelial cells, we speculated that the downregulated hub genes would provide more information about the influence of vessels in the VBCTs. The four downregulated hub genes were determined namely BMP4, C3 protein, CXCL13, and DKK1. BMP4 and C3 were significantly expressed in normal tissue as compared to cancerous breast cancer tissues **(Figure 5d, e)** whereas CXCL13 and DKK1 is observed to be expressed in breast cancer tissue as compared to normal breast tissue **(Figure 5f, g**).

## 3. Discussion

The theory that there is already a prior interaction between primary tumor and secondary site before metastasis is not new.^42^ For instance, previous reports have shown that lung tumors can remotely activate the OBs, and the OBs, in turn, supply the lung tumors with tumor-promoting neutrophils even before the inception of metastasis.^43^ Another *in-vivo* study showed that OBs-derived hypoxia-inducible factor led to distant breast cancer growth and dissemination, partly due to the enhanced production of CXCL12.^19^ The study also showed that osteoclasts did not possess a similar ability to regulate distant BCCs, which highlights the importance of OBs in influencing primary tumor site.^19^ A recent article showed that colorectal cancers release extracellular vesicles that activate fibroblasts in distant organs.^44^ The fibroblasts then produces inflammatory cytokines that promote primary tumor growth.^44^ Therefore, it is evident that the pre-metastatic indirect interaction between secondary and primary sites could illuminate several hidden yet important pathways that initiate the metastasis process.

In human bone, OBs account for 4-6% of the total bone cells and deposit bone matrix.^45^ OBs maintain a homeostatic bone environment together with osteoclasts, the bone-degrading cells and resident osteocytes.^45^ Breast cancer metastasis to the bone disrupts this homeostasis and leads either to enhanced bone deposition (osteoblastic) or enhanced bone resorption (osteolytic). Several studies in the past have already elucidated the role of BCCs in the fate of OBs.^46, 47^ These studies illuminate how the BCCs remotely prime the secondary bone sites before or after homing. However, the role of the secondary bone site on the primary tumor prior to metastasis hasn’t been studied in its entirety.

Cancer cells not only possess the intrinsic ability to proliferate and metastasize but also rely on their surrounding tumor-microenvironment for their progression. The TME is a crowded milieu comprising of multiple cell types such as fibroblasts, endothelial cells, and ECM moieties that constantly interact with one another to modulate tumor fate. The scaffold-free VBCTs that we fabricated are composed of the cancer cells at an interface of vasculature and connective tissue and provide a realistic alternative to conventional 3-D models. Our results indicate that the presence of MDA-MB231 in VBCT resulted in sectioned and abnormal vessels and this abnormality was accentuated in the presence of OBs.

Solid tumors require vessels for growth and metastasis and therefore enforce the production of more BVs. However, cancer cells not only rely on angiogenesis to initiate the metastatic process but can also hijack pre-existing vessels in the tumor microenvironment in a non-angiogenic process called vessel co-option.^48, 49^ Cancer cells, during rapid proliferation in the tissue, also compress the surrounding vasculature in the microenvironment thereby leading to vessel collapse.^50^ Recently, it was shown in an *in-vitro* 3-D model that cancer cells’ interaction with the vasculature led to compression and destruction of the vessel.^51^ In another study, it was shown that pancreatic ductal carcinoma cell (PDAC) ablates endothelial cells both in *in-vitro* and *in-vivo* settings that lead to hypo-vascularization in the PDAC tumor tissue.^52^

In a confined tumor-microenvironment, cells compete for space and nutrients. Cancer cells are highly proliferative in comparison to endothelial cells, which could lead to the elimination of the ‘weaker’ cell population (endothelial cells) by the ‘fitter’ cell population (MDA-MB231).^52^ The scaffold-free model fabricated in a confined cell-culture insert area includes three different cell types in large numbers, each competing for the available nutrients and space, which could explain the elimination of endothelial cells. Furthermore, it could be hypothesized that the presence of OBs and their secreted factors such as CXCL12 could lead to enhanced cellular proliferation and therefore further lead to reduced availability of the limited nutrient to individual cells. We observed via cell proliferation XTT assay that CXCL12 leads to a slight increase in MDA-MB231 and HUVECs proliferation, and a significant increase in proliferation of fibroblasts **(Figure S1h).**

Several other factors could affect angiogenesis and vascular fate. One such factor is the progression stage of the tumor. For instance, it has been shown previously in gliomas that the vessels undergo three stages of development: a) tumor-vessel interaction phase b) transition phase where vessel apoptosis takes place followed by c) angiogenic phase.^53^ The increase in VEGF and the presence of thin vascular sprouts in VBCT suggest a transition phase preceding angiogenesis. For these phases to occur, tumor cells have to be in close proximity to the vessels since these phases require direct cell-cell interactions. The advantage of our model is the ability of cancer cells to directly interact with BVs **(Figure S3)**. Lastly, the influence of the highly proliferating TNBC on vasculogenesis (apart from angiogenesis) should also be taken into consideration since MDA-MB231 cells are mixed together with endothelial cells and not introduced to fully-formed vessels. Future experiments where MDA-MB231 cells are introduced after the formation of BVs in the model could elucidate different angiongenic mechanisms.

We show a number of interesting upregulated hub genes from the gene array data between NHMO and NHM samples where we studied the effect of OBs on VBCTs. We observed HSP90N as an upregulated hub gene that has been previously linked to aggressiveness in breast tumors.^54, 55^ HSP90N inhibitors have been used in the past for cancer therapy and could also have an implication in bone metabolism.^56^ Interestingly, HSP90N expression is also associated with enhanced bone metastasis from prostate and breast tumors.^57, 58^

The other hub genes EGFR, RPS27A and CYCS have also been previously identified as hub genes in TNBC extracted from several gene expression datasets.^59^ The overexpression of EGFR, another upregulated hub gene, leads to poor prognosis in breast cancer patients.^60, 61^ EGFR results in cancer cell proliferation, aggressiveness and metastasis.^62^ Interestingly, expression of EGFR has also be linked to increased metastasis of cancer cells to bone.^62^ For instance, previous results have shown that EGFR signaling can lead to osteoclastogenesis by influencing OPG and MCP1 signals in osteoblasts.^63^

The changes in gene expression between NHMO and NMO were studied to deliberate the impact of BVs. We hypothesized the presence or absence of BVs would lead to modified gene expression. DKK1, an antagonist of the Wnt/β-catenin signaling pathway was detected as a downregulated hub gene. Previously, it has been shown that DKK1 functions as a tumor suppressor.^64^ Interestingly, upregulation of DKK1 has shown to be involved in specific breast cancer metastasis to bone.^65, 66^ Furthermore, tumor-derived DKK1 also dictates the OBs fate by inhibiting OBs differentiation and facilitates osteolysis.^67^ It is intriguing that in the presence of BVs in the cancerous tissue, DKK1 was observed as a downregulated hub gene. In malignant gliomas, it was previously reported that hypoxia led to an upregulation of DKK1 as compared to normoxia.^68^ It could be theorized that in the absence of BVs and their secreted growth factors in the non-vascularized tissue, DKK1 was upregulated which in turn could result in the enhanced preference of breast cancer metastasis to bone. A more thorough future investigation into the role of BVs in DKK1 expression could highlight many possible mechanisms in bone metastasis. Another downregulated hub gene BMP4 has previously shown to inhibit proliferation but accentuate cell migration in breast cancer.^69^ BMP4 can act as both tumor suppressor and promoter.^70^ One study showed that BMP4 upregulation is associated with reduced metastasis and the downregulation of BMP4 has been associated with more aggressive breast cancer.^71^ Another study showed that mice treated with BMP4 showed a positive correlation with bone metastasis.^72^ Finally, downregulated hub genes complement protein C3 and CXCL13 both could have implications in breast cancer progression and modulation in bone microenvironment.^73, 74^

Together, the developed *in-vitro* model could is suitable to study several pathways involved in the progression of metastasis and organotropism. In this study, we show that indirect culture with OBs leads to a change in the gene expression profile of VBCTs. This emphasizes the point that reciprocal OBs secreted factors could be an important parameter in early metastasis events in breast cancer. Future work would include other main cellular players such as osteoclasts and a 3-D bone microenvironment together with functional immune cells in the VBCTs to elucidate the role of different cell types in cancer progression or dormancy.

## 4. Conclusion

In summary, we present a 3-D co-culture system utilized to understand the interaction between several cell types involved in breast cancer metastasis to the bone. The fabricated VBCTs provide a realistic microenvironment to the TNBC cells with the presence of fully formed vasculature, surrounding fibroblasts and cell-secreted ECM. The simplicity of the culturing and harvesting of the 3-D model allowed a gene-level study of the influence of OBs and BVs on the engineered cancer tissue. These results provide a proof-of-concept for the utilization of such models for the understanding of time-dependent metastatic progression. Such models in the future can be used in predicting essential biomarkers for diagnosis and for pre-clinical testing of novel therapeutic agents.

## 4. Methods and Materials

### 4.1 Cell culture and media condition

GFP^+^ MDA-MB231 cells were kindly provided by Dr. Julia Steitz, Institute of Laboratory Animal Science, RWTH University Hospital, Aachen, Germany. The cells were cultured in DMEM (11965092, Thermofisher Scientific) with 10% fetal bovine serum (FBS). Osteoblasts were procured from Sigma Aldrich (406-05F), and cultured in Osteoblast growth media (41-7500, Sigma Aldrich). Fibroblasts were obtained by isolation of skin biopsies that were surgically extracted from healthy volunteers (Department of Dermatology, RWTH Aachen University Hospital, Germany). The extraction and isolation were conducted in accordance with the Declaration of Helsinki principles and was approved by the ethical committee of RWTH Aachen University Hospital, Germany. Fibroblasts were cultured in DMEM with 10% FBS. HUVECs (Lonza, Passage <4) were cultured in an endothelial growth medium (C-22111, Promocell). For the culture of VBCTs, DMEM: EGM (1:1) with 10% fetal bovine serum (FBS, Biowest) and 50 μg/ml Ascorbic acid (A92902, Sigma) was used.

### 4.2 Migration assay

OBs (passage<5) were cultured in 24 well plates (7×10^4^ cells) and allowed to cultivate for a day. MDA-MB231 cells were cultivated in serum-free DMEM media for 24-hours. Required number of cells was treated with 25 μg/ml AMB3100 (EMD Millipore) for 1 hour. Cell culture inserts (8μm pore size, Corning) were pre-treated with DMEM medium for 2 hours prior to cell addition. 1×10^4^ treated or untreated MDA-MB231 cells were added to the upper compartment of the 8μm pore size cell culture inserts in 250 μl of serum-free DMEM (with 0.1%BSA) whereas 600 μl of serum-free DMEM (with 0.1% BSA in media) was added to the lower compartment. After 1 hour of attachment time, inserts were transferred into the well plate containing OBs and fresh media was added. CXCL12 (Sigma, SRP3276) in serum-free DMEM media was added to appropriate chambers depending on the experimental requirement. After 18 hours of the assay, the lower compartment was stained with DAPI and images were taken in a fluorescent microscope (10X magnification). Three separate regions were selected per insert. Experiments were done in triplicates.

### 4.3 XTT assay

5×10^4^ fibroblasts, HUVECs or MDA-MB231 cells were seeded on 96-well plates. In each cell conditions, 0 ng/ml (vehicle: DMSO), 50 ng/ml and 100 ng/ml of CXCL12 were added for 24 hours in respective serum-free media (0.1% BSA). XTT (11465015001, Sigma) was utilized as per the manufacturer’s protocol. The specific absorbance at 460 nm was observed and the unspecific absorbance at 630 nm was subtracted.

### 4.4 ECM-coating and accumulation to fabricate cancer models

Fibroblasts were coated layer-by-layer using ECM coating and accumulation technique as previously described.^26^ Briefly, fibroblasts were exposed to 0.04 mg/ml FN (sc-29011A, SCBT) and 0.04 mg/ml G (G9391, Sigma) solutions with an intermediary PBS washing step. After 9 steps of alternate exposure to FN and G, the coated cells were mixed with HUVECs in 15:2 ratios and 2.5×10^4^ MDA-MB231 were mixed with the HUVECs and coated fibroblasts. The models were cultured for 24 hours in 0.4 μm pore sized inserts (3470, Corning) and the inserts were transferred to well plates containing OBs. Prior to the fabrication of models, 7×10^4^ OBs were seeded on 24-well plates and cultured for 2 days.

### 4.5 Immunostaining of tissues

Tissues were extracted out of the cell culture insert, washed twice with PBS. The harvested tissues were fixed in 4% PFA for 30 minutes at room temperature (RT). The tissues were washed and suspended in 0.1% Triton-X 100 for 10 minutes. The unspecific antibody binding was blocked by suspending the tissue in 3% BSA (Sigma, Germany) in PBS for 1.5 hours in RT. After PBS wash (2X), mouse anti-human CD-31 primary antibody (MA5-13188, Thermofischer Scientific, 1:100 dilution in PBS) was added to the tissue and incubated overnight in 4°C. After 24 hours, the tissues were washed with PBS (3X, 5 minute interval in the shaker) and goat anti-mouse Alexa 594 (A11005, Invitrogen) 1:200 dilutions was added for 3 hours in RT. Tissues were washed with PBS (3X, 5 minute interval in shaker). The tissues or 2-D MDA-MB231 were further stained with DAPI (1:1000, 5 minute incubation in RT) if required. Finally, the samples were prepared for confocal microscopy (Leica TCS SP8).

### 4.6 Microscopy and image analysis

Stained VBCTs were extracted from the cell culture inserts, fixed on a glass slide and investigated under the confocal microscope. Samples were observed for vessels (CD31) using an excitation wavelength of 594nm (red), GFP^+^ MDA-MB231 using an excitation wavelength of 488nm (green) and DAPI (nucleus) with an excitation wavelength of 405nm. Images were taken at a resolution of 1024×1024 pixels using either 10X dry objective, 20X Oil immersion objectives or 63X Oil immersion objectives depending on the experimental requirements. Vessels were analyzed for vascular density, area percentage and junction point using AngioTool as per software instruction.^75^ All illustrations in the paper were made using Biorender.com

### 4.7 ELISA

The cancer models were cultivated with and without OBs for 7 days with alternative day media changes. Media was harvested and centrifuged (1500 RPM, 10 minutes, 4°C) and the supernatant was collected. The Human VEGF165 Standard ABTS ELISA Development Kit (900-K10; Peprotech) and ABTS ELISA Buffer Kit (900-K00; Peprotech) were used as per manufacturer’s protocol.

### 4.8 Gene chip microarray

To generate amplified sense-strand cDNA, 300 μg purified mRNA of each model were processed using the WT Expression Kit (Ambion, Austin, TX, USA). Affymetrix® GeneChip® WT Terminal Labeling Kit Assay (Affymetrix, Inc., Santa Clara, CA, USA) was used for fragmentation and labeling. Prepared hybridization cocktails were applied to Clariom™ S assay (Thermo Fisher Scientific). The assays were washed and stained using the GeneChip® Fluidics Station 450 (Fluidics Protocol FS450_0001). Arrays were scanned using Affymetrix GeneChip^®^ Scanner 3000 7G controlled by GeneChip^®^ Operating Software (GCOS) version 1.4 to produce CEL intensity files.

### 4.9 Gene analysis and gene ontology

The raw CEL files were analyzed using TAC (Applied Biosystems) and subsequent normalization was performed as per software instruction. The expression values were observed and the upregulated genes with fold change ≥2 and P value<0.05 and downregulated genes with fold change ≤−2 and P value<0.05 were chosen for further analysis. The gene ontology of the acquired DEGs was performed in Metascape (http://metascape.org). ^76^ The DEGs were fed into the online system and the pathway analysis was conducted utilizing a combination of GO Biological Processes, KEGG Pathway, Reactome Gene Sets and WikiPathways. Ontology terms with a p-value < 0.01, a minimum count of 3, and an enrichment factor > 1.5 were considered. The volcano plots were constructed using VolcaNoseR.^77^

### 4.10 PPI network analysis and prediction of hub genes

The entire upregulated or downregulated genes were fed into STRING online application that predicts the PPI network.^78^ A medium confidence score of 0.4 was selected. The formed PPI network was uploaded to the Cytoscape software (version 3.8.1).^79^ The hub genes from the entire network were obtained by the Cytohubba application.^80^ The top nodes were ranked utilizing the MNC network scoring method.

### 4.11 Obtaining highly interconnected clusters

To obtain the clusters, the genes were fed to the MCODE app in the Cytoscape software.^81^ The degree cut-off was selected as 2, node cut-off score as 0.2 and K-score as 2. The clusters with a MCODE score greater than 10 was selected. The functional enrichment analysis was conducted in Cytoscape where Reactome pathways were selected.

### 4.12 Expression analysis and survival plots

The gene expression analysis was performed in GEPIA2 with p-value cut-off of 0.05.^82^ BRCA and Basal-like/Triple negative datasets were selected. The disease free survival (DFS) was analyzed for the hub genes using media cut-off (cut-off high 50%, cut-off low 50%) in GEPIA2.^82^ BRCA and Basal-like/Triple negative datasets were selected for plotting the Kaplan Meier plot.

### 4.13 Statistics

Statistical analyses were performed using GraphPad Prism 9.0.0 software. All experiments were performed in triplicates. Data are presented as mean ± standard deviation unless otherwise stated. Data are analyzed using one-way ANOVA with Tukey’s multiple comparison test. Significance was considered when p < 0.05. Individual p-values are stated in the graph.

## Acknowledgments

This work was supported by the Deutsche Forschungsgemeinschaft (DFG) in the framework of the Research Training Group “Tumor-targeted Drug Delivery” grant 331065168. Microarray analysis was supported by the Genomic Facility, a core facility of the Interdisciplinary Center for Clinical Research (IZKF) Aachen within the Faculty of Medicine at RWTH Aachen University. The authors thank Dr. Sebastian Huth for the scientific discussion. We owe special acknowledgment to the cooperation of Prof. Jens Baron, Department of Dermatology, RWTH University Hospital, Aachen, Germany

## Supporting Information

**Figure S1.**
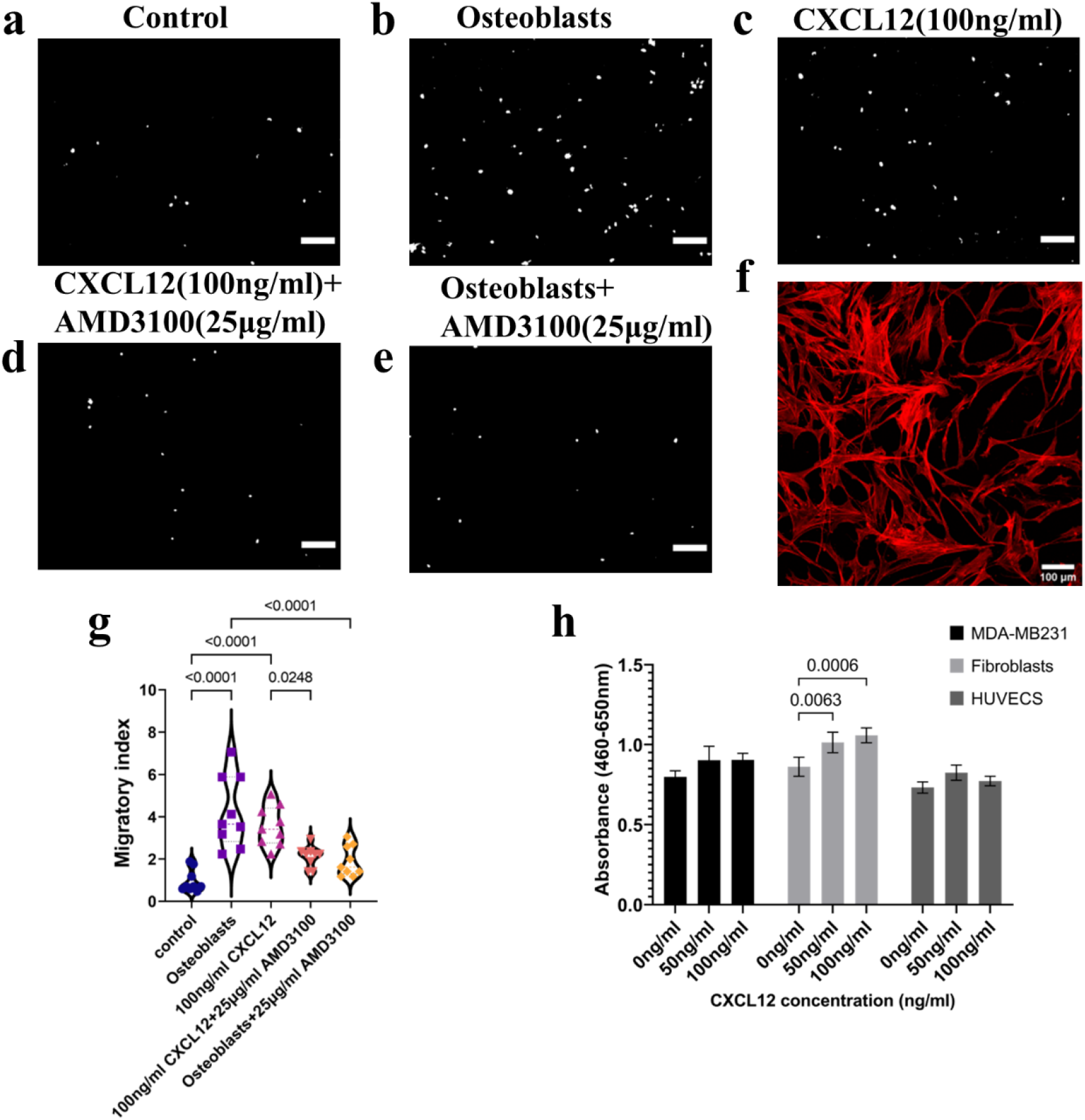
Migration of MDA-MB231 in a) Serum free media (control), b) Presence of osteoblasts in the well-plate in serum-free media c) Presence of 100 ng/ml of CXCL12 at the bottom of the insert in serum-free media d) Pre-treated MDA-MB231 with 25 μg/ml AMD3100 slower migration towards CXCL12 e) Pre-treated MDA-MB231 with 25 μg/ml AMD3100 slower migration towards osteoblasts, scale bar=100 μm f) morphology of OBs on the well plate, scale bar=100 μm g) significant difference in migration of MDA-MB231 in media vs osteoblasts and media vs CXCL12 added in the bottom, significant difference in migration observed after pre-treatment with AMD3100 towards osteoblasts and CXCL12 (n=3, with 3 different regions of the image) h) XTT assay shows slight increase in absorbance values in BCCs and HUVECs treated with 50 ng/ml and 100 ng/ml of CXCL12 and significant increase in Fibroblasts treated with 50 ng/ml and 100 ng/ml CXCL12 (n=3). Statistics used was ordinary one-way Anova with Tukey test.

**Figure S2.**
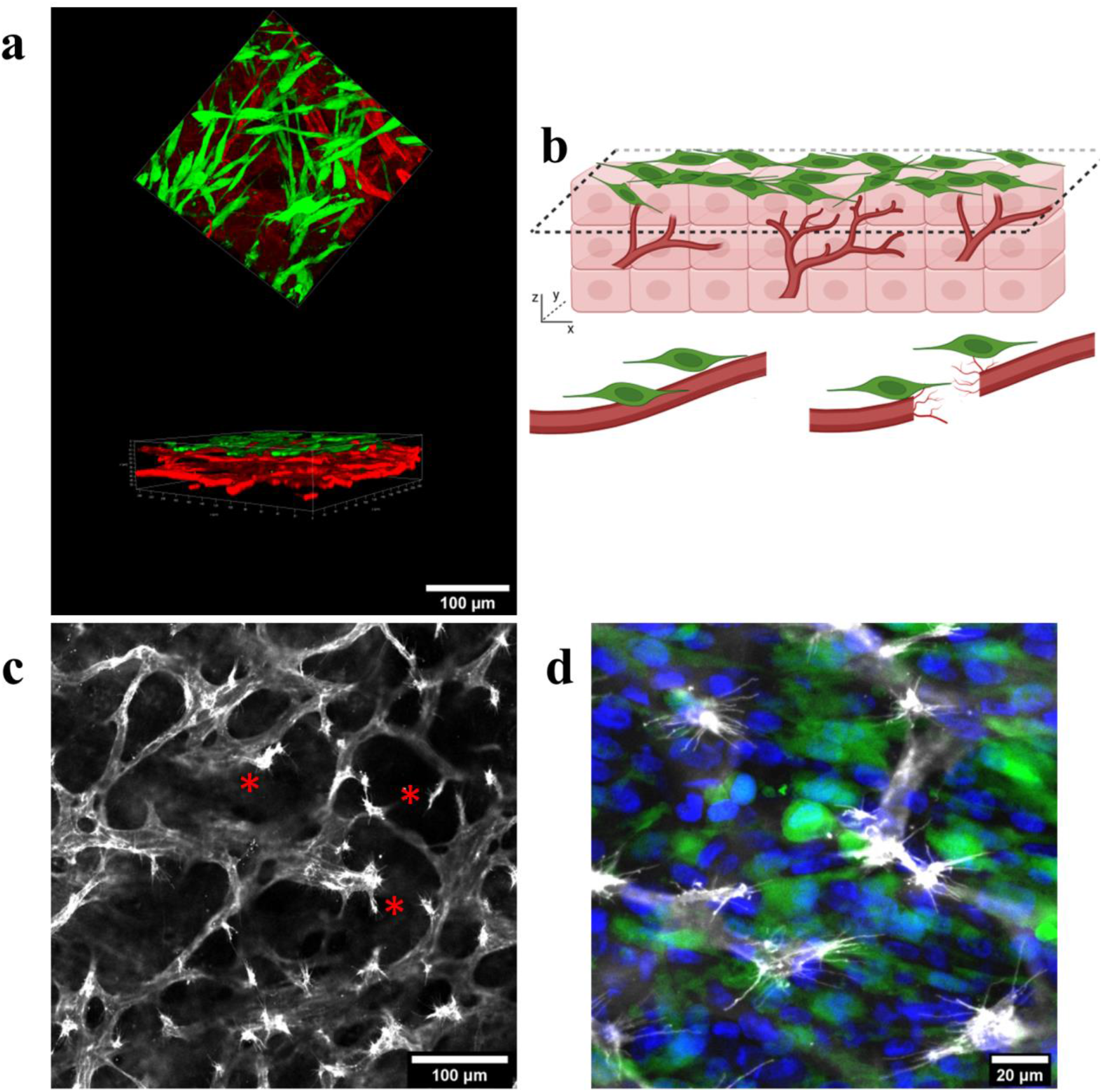
a) Top view and side view of MDA-MB231(green) seeded on top of vascularized tissue (BVs= PECAM-1 stained red), scale=100 μm b) schematic illustration of the fate of vessels in direct contact with BCCs c) confocal images show broken vascular endpoints when in contact with MDA-MB231 cells, scale=100 μm d) magnified image shows the endpoints with sprouts (vessels= PECAM-1 stained, false colored gray, MDA-MB231=green), scale bar=20 μm

**Figure S3.**
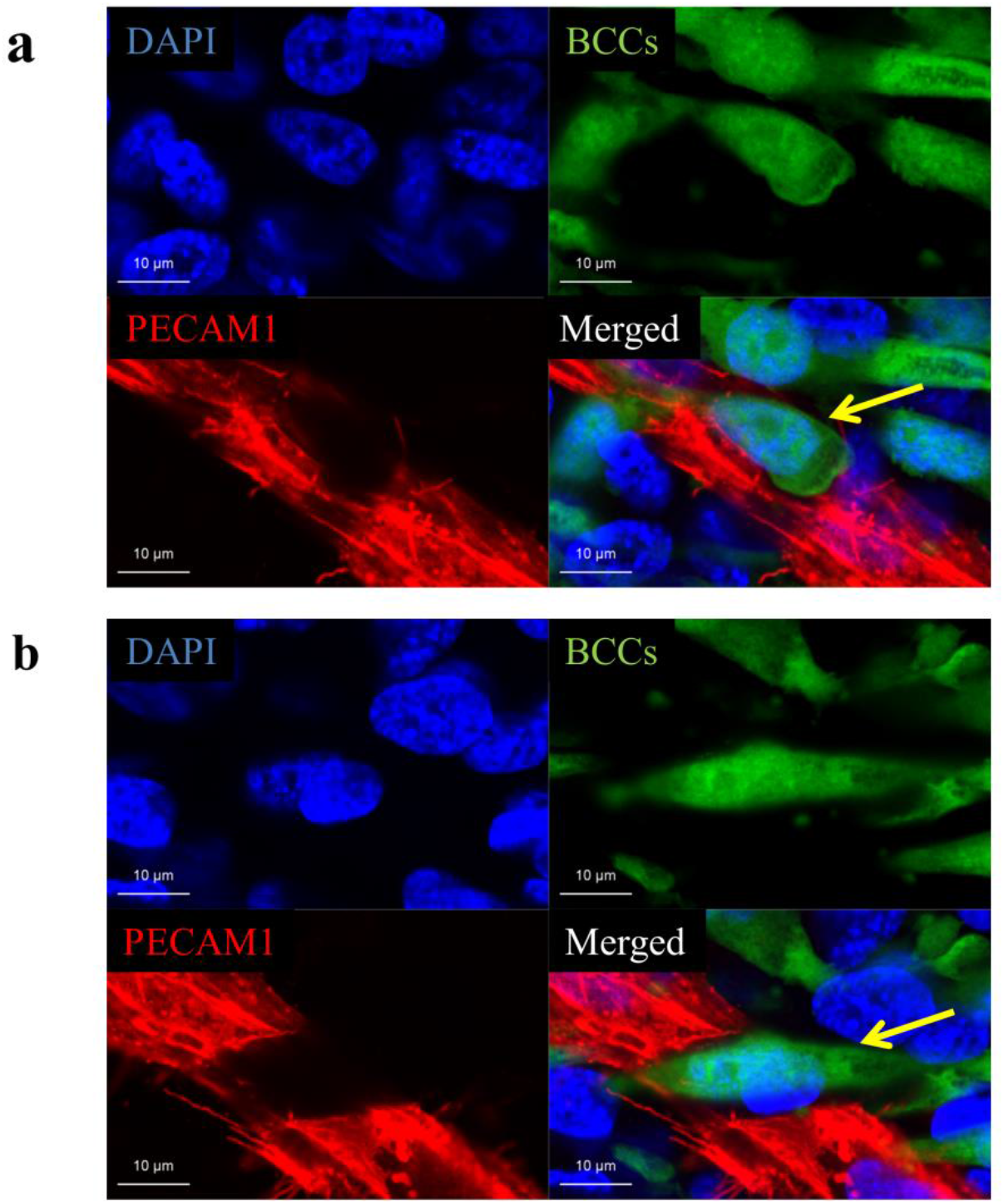
a) Direct cell-cell contact between BCCs and vessels (DAPI=blue, BCCs=green, PECAM-1=red), scale bar=10μm b) BCCs penetrates and sections the vessels during direct contact, scale bar=10 μm

**Figure S4.**
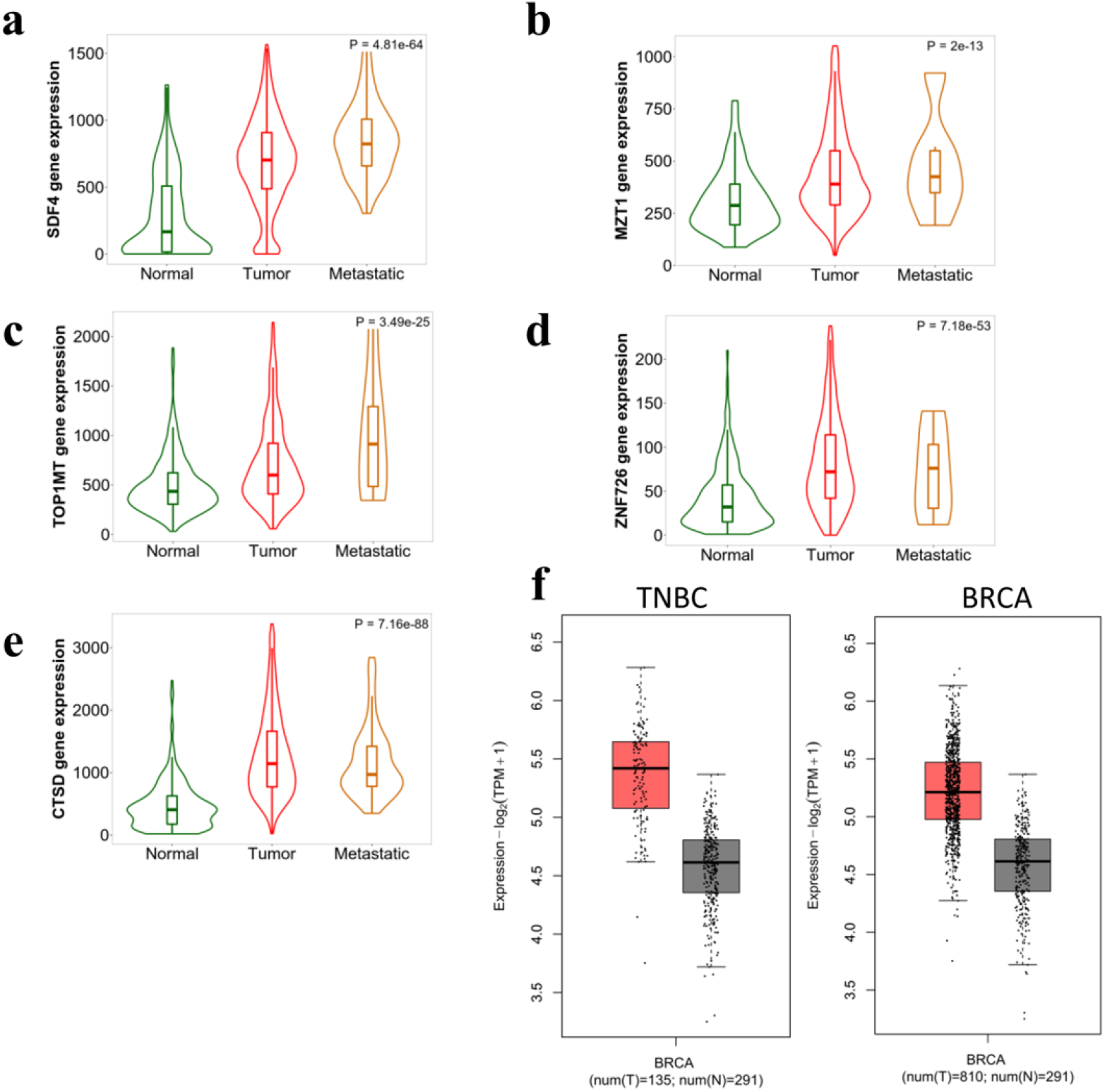
Expression of the top five upregulated genes (NHMO vs NHM) in normal, tumor and metastatic tissue patient database (TNM plotter). Expression analysis of a) SDF4, b) MZT1, c) TOP1MT, d) ZNF726, and e) CTSD show higher expression of the genes in metastatic BRCA tissues as compared to normal breast tissues. f) The gene signature comprising of the top five upregulated genes were fed to the GEPIA2 online software. The gene signature is observed to be expressed more in specific TNBC tissue and overall BRCA tissue as compared to normal tissue.

**Figure S5.**
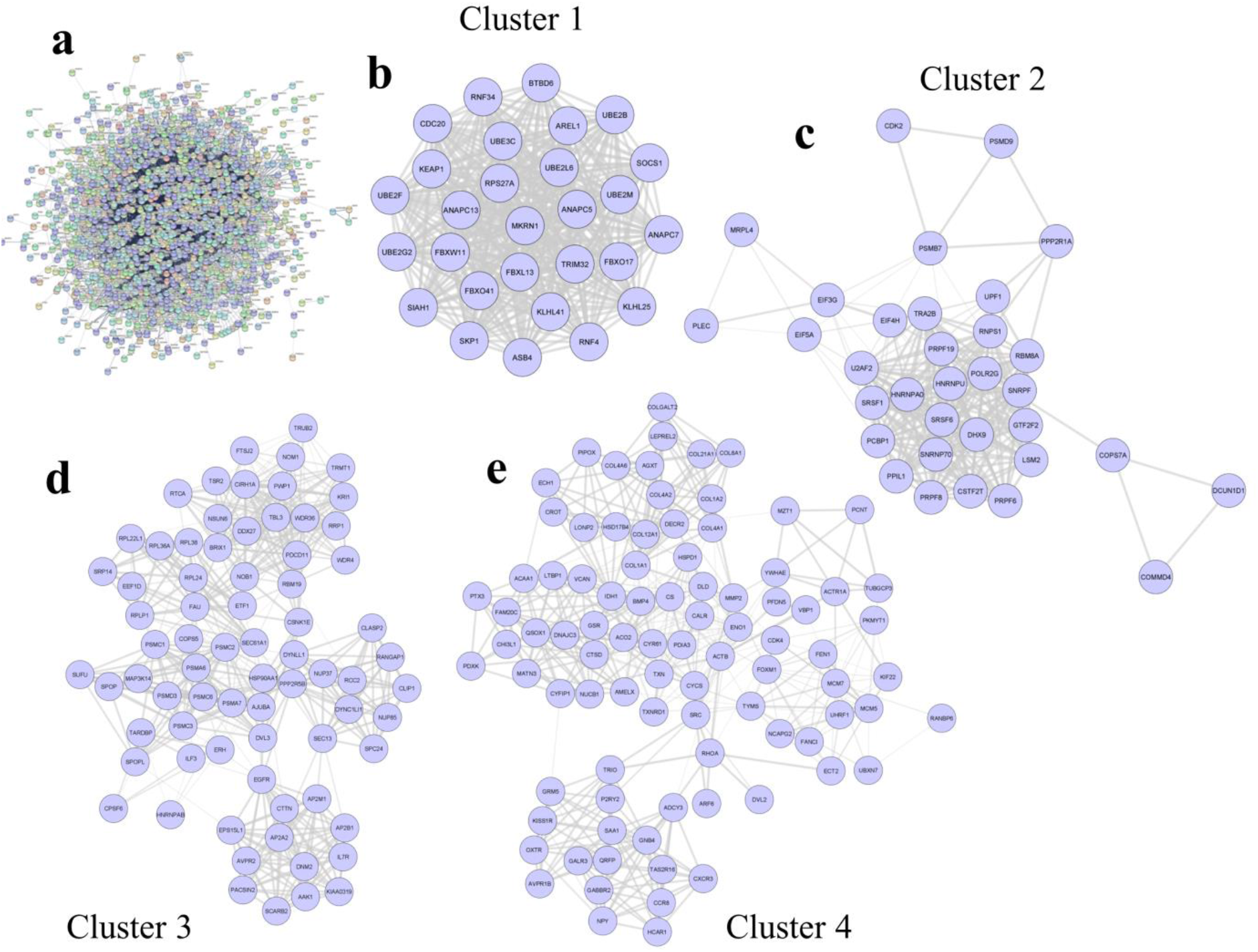
Upregulated protein clusters in NHMO vs NHM a) Overall upregulated protein-protein interaction network b, c, d, e) Four highly upregulated upregulated clusters (Clusters 1, 2, 3, and 4) acquired from MCODE application in the Cytoscape software.

**Table S1.**
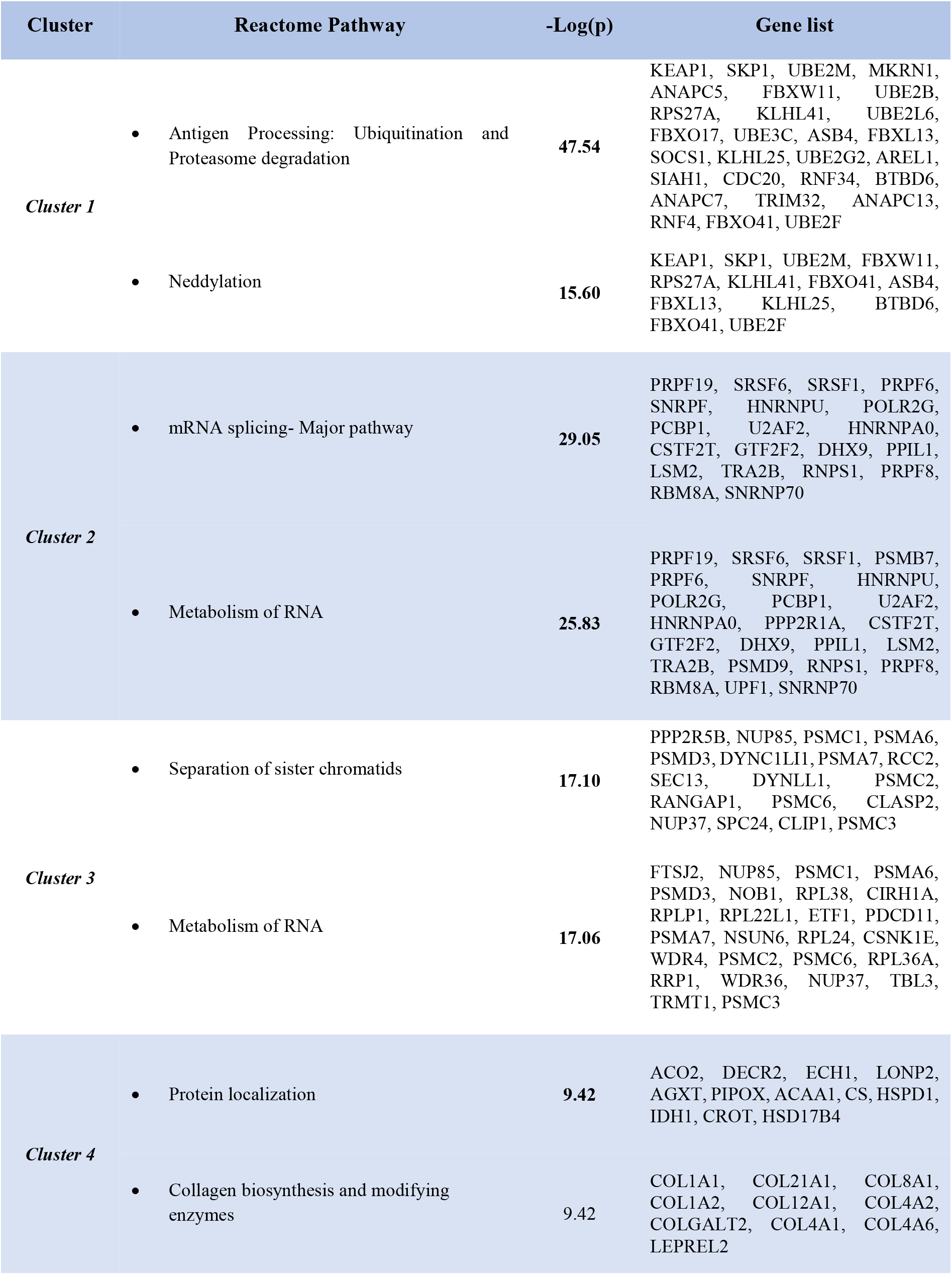
shows the highly interconnected upregulated clusters (NHMO vs. NHM) and the genes involved in each cluster.

**Figure S6.**
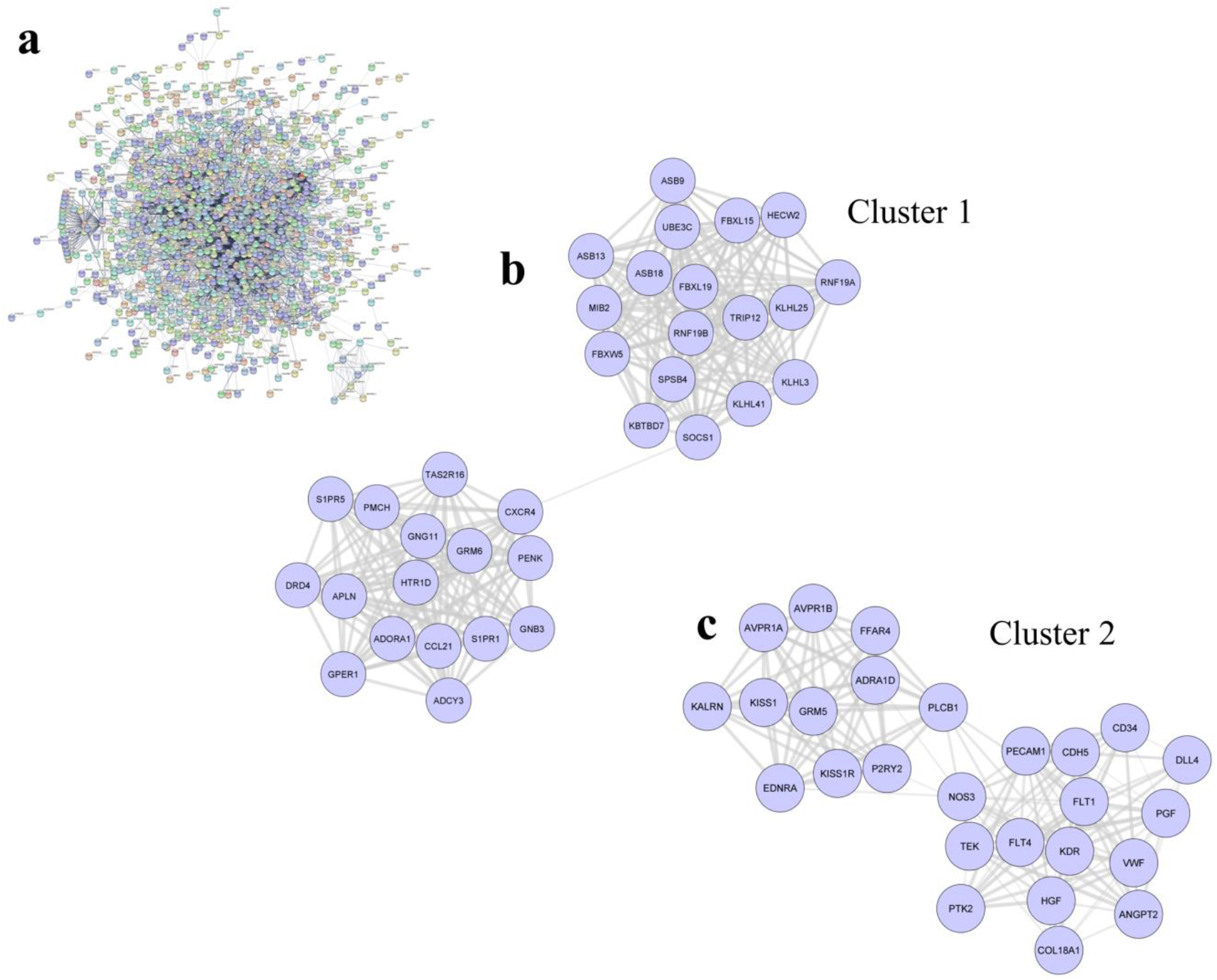
Upregulated protein clusters in NHMO vs NMO a) Overall upregulated - interaction network, b, c) Two highly interconnected upregulated clusters (clusters 1 and 2) acquired from MCODE application in the Cytoscape software.

**Table S2.** Upregulated gene ontology and gene symbols (NHMO vs NMO)

| Category | Term | -Log(p) | Gene symbols |
| --- | --- | --- | --- |
| GO:0030855 | • Epithelial cell differentiation | 7.702 | BMP2,CD34,CDH5,CDSN,COL4A1,CYP1A1,DAB2,DLX3,S1PR1,EVPL,FLNA,NR5A2,B4GALT1,ID1,ID2,ID3,IL1B,JAG2,JUP,KDR,KRT5,KRT9,KRT15,KRT86,LORICRIN,MEF2C,AFDN,PECAM1,PODXL,POU4F3,PRKCH,PTK6,S100A7,SOX4,SPRR2G,SPRR3,ST14,TBX1,TCF15,TIE1,TJP1,CXCR4,PKP4,KRT75,TGM5,SLC4A7,RAPGEF2,MTSS1,CCNO,KLF2,UPK1A,PLCB1,HEY1,SCRIB,MAFF,PPP1R16B,ABCA12,LCE2B,F11R,SOX18,COL18A1,KRTAP4-5,IL31RA,KRT72,KRTAP15-1,KRTAP6-1,KRTAP20-1,KRT79,LIPM,LCE1C,LCE1D,KRTAP12-1,KRTAP10-6,KRTAP10-5,KRTAP5-4,MIR10A,KRTAP5-7,CCDC88C,LIPN,COL1A2,COMP,INHBA,ITGA6,SRF,AP3B1,HLA-DRB1,,KRTAP21-1, KRTAP29-1 |
| GO:0006935 | • Chemotaxis | 5.295 | ADAM8,ANGPT2,CDH13,LYST,CXADR,S1PR1,EFNA1,EGR2,EPHA3,EPHA4,EPHB1,FOXG1,FLT1,HGF,IL6R,KDR,KIF5A,LAMA2,MST1,NCAM1,NOTCH3,OR1D2,PDE4B,PGF,PIK3CB,PIK3CG,POU4F3,PPIB,PTK2,S100A7,CCL14,CCL15,CCL21,SEMA3F,SLC12A2,SPN,CXCR4,SEMA3B,PLA2G10,KALRN,DOCK4,SEMA6B,SEMA4B,DPYSL4,B3GNT2,PLXND1,SCRIB,DAPK2,PRKD2,ARHGEF16,LEF1,SEMA5B,LTB4R2,DPYSL5,PLEKHG5,MTUS1,PREX1,MMP28,SHANK3,TIRAP,EMB,CMTM2,UNC5B,DEFB109B,ECSCR,FLNA,ID1,TACSTD2,MEF2C,MYH9,NPTX1,SRF,ST14,CDKL5,TBCD,TEK,USP9X,FZD4,RAPGEF2,KIAA0319,PLPPR4,WDR1,NBEAL2,COBL,PPP1R9A,RCC2,HECW2,TNN,COL18A1,FERM T3,ACTBL2,CCDC88C,SYT1,UGT8,RASAL1,RIMS1,SYT17,CPNE5,CACNG7,NES,ABLIM1,RGS3,RNASE1,ABLIM2,MYL12B,FKBP1B |
| GO:0030335 | • Positive regulation of cell migration | 5.169 | ADAM8,BMP2,CDH5,CDH13,D<br>AB2,S1PR1,ETS1,FLNA,FLT1,F<br>LT4,GPER1,HGF,HYAL1,IL1B,I<br>L6R,ITGA6,ITGA2B,KDR,MCA<br>M,MMP14,NOS3,PECAM1,PGF,<br>PIK3CB,PIK3CG,PODXL,PTK2,<br>S100A7,CCL21,SEMA3F,SPN,T<br>EK,TJP1,CXCR4,SEMA3B,RAP<br>GEF2,DOCK4,HDAC9,SEMA6B<br>,SEMA4B,PLK2,CLASP2,DAPK<br>2,SUN2,PRKD2,NOX4,LEF1,SE<br>MA5B,PREX1,DOCK8,PLVAP,<br>CD99L2,FERMT3,TIRAP,MIR1<br>0A,SMIM22,FAM110C,MIR939,<br>RNASE10,CHMP4C,ANGPT2,M<br>ST1,MTUS1,MMP28 |
| hsa05200 | • Pathways in cancer | 5.009 | ADCY3,AREG,BAK1,BMP2,CA<br>SP7,CCND3,COL4A1,CSF2RB,<br>E2F1,EDNRA,EPAS1,EPOR,ET<br>S1,MECOM,FGF11,FLT4,GNA<br>Q,GNB3,GNG11,MSH6,HGF,IF<br>NA2,IFNA5,IFNAR1,IL2RA,IL3<br>RA,IL6R,IL7,IL12B,ITGA6,ITG<br>A2B,JAG2,JUP,LAMA2,MMP1,<br>NOTCH3,PGF,PIK3CB,PMAIP1,<br>PTK2,RAD51,RNASE1,RXRA,T<br>GFB3,ZBTB17,CXCR4,FZD4,T<br>NFRSF25,LONP1,PIM2,PLCB1,<br>HEY1,DAPK2,RASGRP3,LEF1,<br>DLL4,PLEKHG5,EGLN3,IL23R,<br>BMP3,TNFRSF8,FLT1,GDF10,I<br>L1B,IL1RAP,INHBA,KDR,CCL<br>14,CCL15,CCL21,TNFSF4,TNF<br>SF10,IL32,IL20RB,CRLF2,IL25,<br>RELT,IL31RA,CCL15-CCL14,IL31,SOCS1 |

**Table S3.**
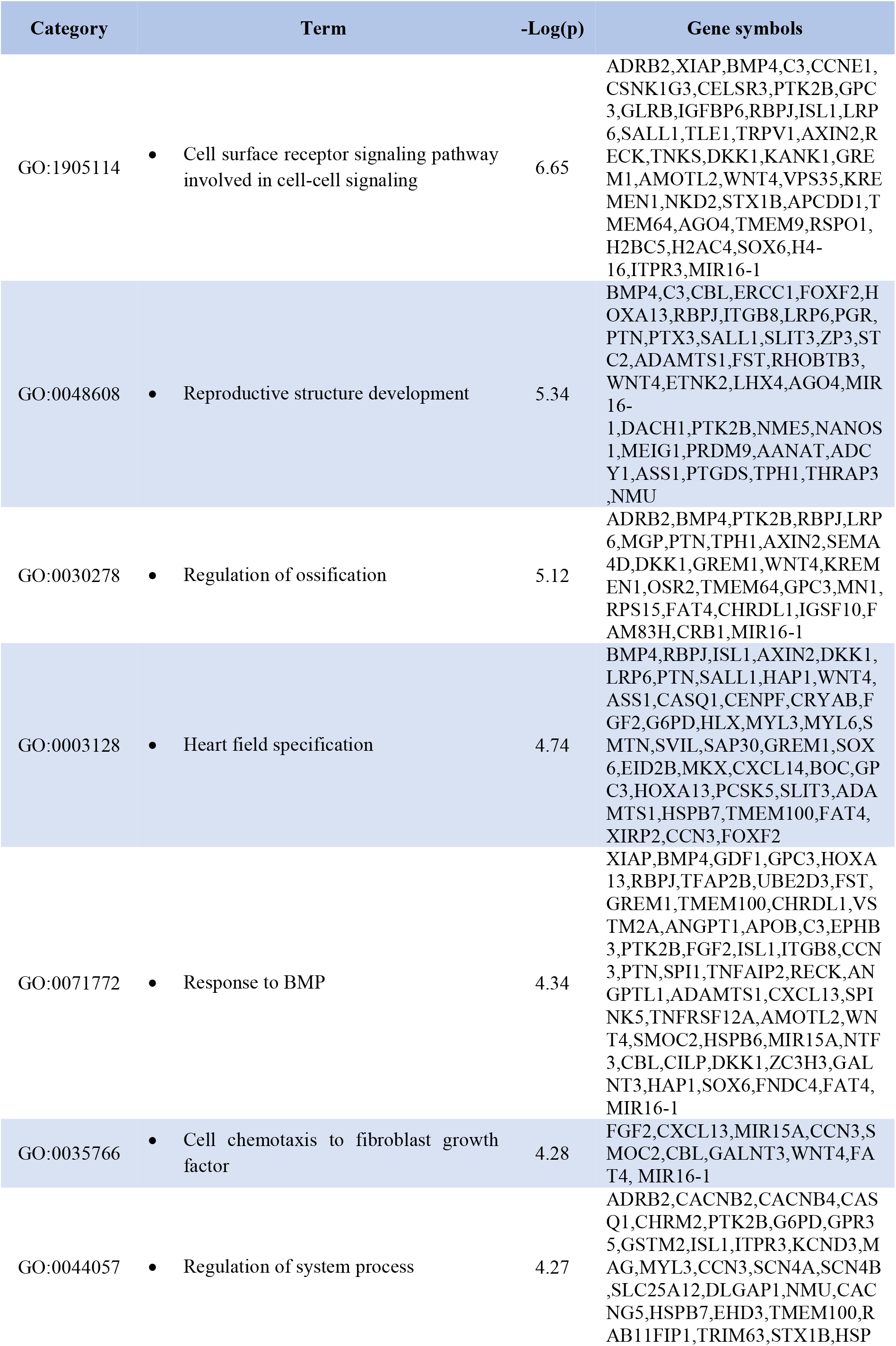
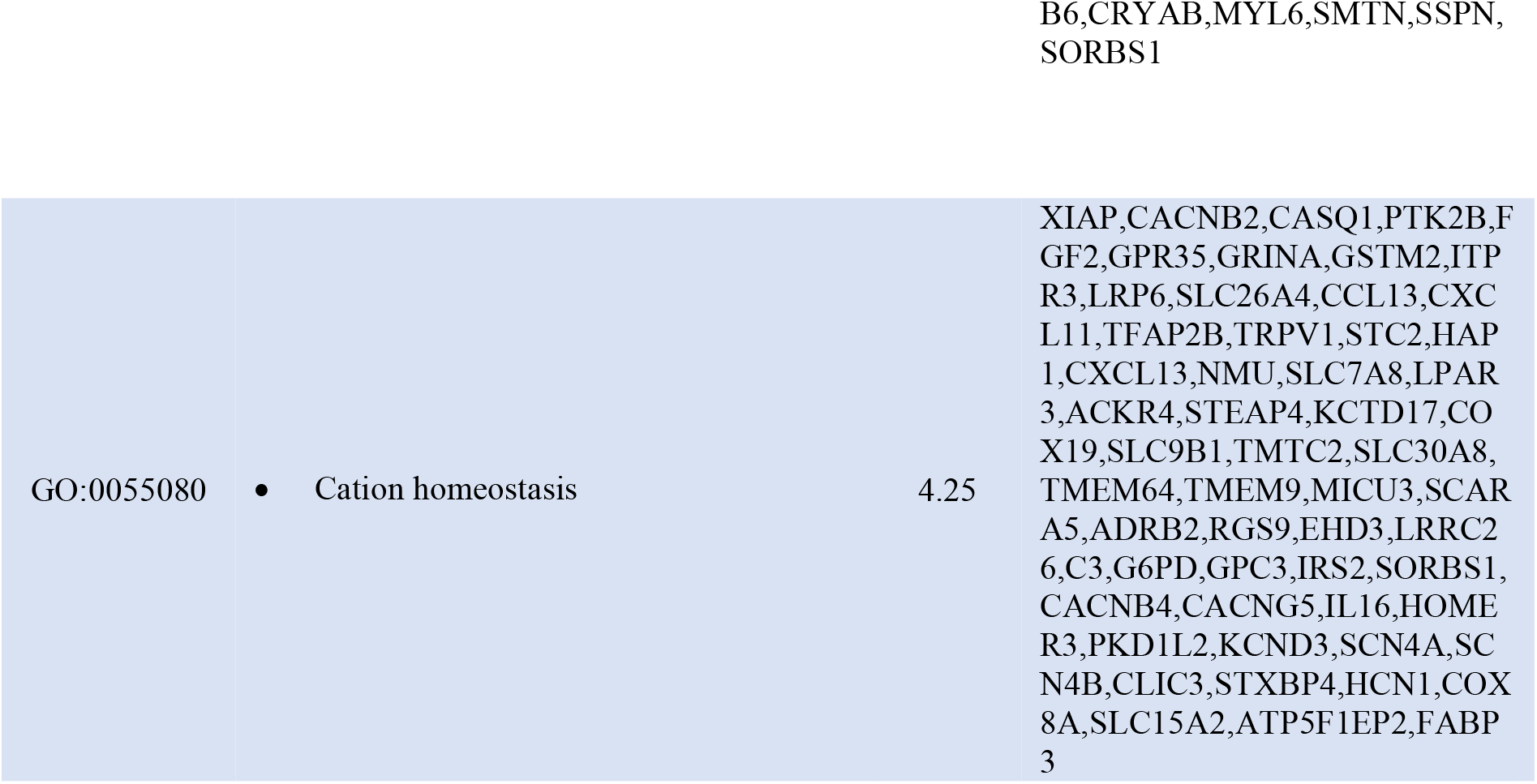
Downregulated gene ontology terms and gene symbols (NHMO vs. NMO)

